# Cryo-EM structure of the octameric pore of *Clostridium perfringens* β-toxin

**DOI:** 10.1101/2021.11.23.469684

**Authors:** Julia Bruggisser, Ioan Iacovache, Samuel C. Musson, Matteo T. Degiacomi, Horst Posthaus, Benoît Zuber

## Abstract

*Clostridium perfringens* is one of the most widely distributed and successful pathogens. It causes multiple severe diseases in animals and humans and produces an impressive arsenal of toxins with pore-forming properties, most of them belonging to the hemolysin-like family of β-pore forming toxins (β-PFTs). One of the most potent toxins produced by *C. perfringens* is β-toxin (CPB). This toxin is the main virulence factor of type C strains and essential for the development of a fatal necrotic enteritis in humans and newborn animals. In the present study, we describe the cryo-electron microscopy (cryo-EM) structure of CPB in styrene maleic acid (SMA) discs, which represents the membrane-inserted pore form, at near atomic resolution. We show that CPB forms a homo-oligomeric pore with eightfold symmetry and similar conformation to the hetero-oligomeric pores of the bi-component leukocidins, with important differences in the receptor binding region and the N-terminal latch domain. Intriguingly, the octameric CPB pore complex contains a second 16-stranded β-barrel protrusion atop of the cap domain that is formed by the N-termini of the eight protomers. We propose that CPB defines a new subclass of the hemolysin-like family of β-PFTs s. In addition, we show that the β-barrel protrusion domain can be changed or modified without affecting the pore forming ability, thus making the pore particularly attractive for macromolecule sensing and nanotechnology. The cryo-EM structure of the octameric pore of CPB will facilitate future developments in both nanotechnology and basic research.

## Introduction

One of the most common and evolutionary conserved bacterial virulence mechanisms is the secretion of protein toxins that disrupt cellular membranes by pore-formation. Such pore-forming toxins (PFTs) are used by the pathogens to invade, survive, and disseminate in their hosts. Despite their large diversity, bacterial PFTs share common features. All are secreted as water-soluble monomers and bind to target cells via membrane receptors. Receptor binding leads to an increase in the local concentration, oligomerization and insertion of a stable pore in the cell membrane. This allows uncontrolled exchanges between the extracellular and intracellular milieus, disturbs cellular homeostasis, and leads to diverse reactions ranging from defense mechanisms to cell death (Dal Peraro & van der Goot, 2016). Because of their nearly universal presence in bacterial pathogens, common structures, and mechanisms used PFTs are promising targets for novel anti-bacterial toxin treatment strategies as alternatives or supplementation to increasingly ineffective antibiotic treatments. Moreover, PFTs have gathered much interest in the scientific community beyond bacterial infections. The nano-sized pores that they form are used for “sensing” biomolecules. Many nanopore applications are based on α-hemolysin (Hla), the prototype hemolysin-like β-PFT secreted by *Staphylococcus aureus* (Kasianowicz *et al*, 1996).

The human and animal pathogen *C. perfringens* causes many severe diseases such as wound infections, septicaemia, food poisoning, enterotoxemia, and enteritis (Kiu & Hall, 2018; Songer, 1996, 2010). The bacterium can produce a large arsenal of exotoxins in particular PFTs that cross the membrane as a β-barrel (β-PFT). Many of them belong to the hemolysin-like family of PFTs (Lacey *et al*, 2019; Mehdizadeh Gohari *et al*, 2015; Popoff, 2014; Popoff & Bouvet, 2009). One of the most potent toxins produced by *C. perfringens* is β-toxin (CPB) (Uzal *et al*, 2014). CPB is secreted by *C. perfringens* type C strains and is essential in the pathogenesis of a lethal necrotic enteritis in many animal species and humans (Uzal *et al*., 2014). Based on sequence homology between mature CPB and other bacterial toxins (Supplementary Fig. 1 and Supplementary Table 1), the toxin is a member of the hemolysin-like family of β-PFTs. Within this group, CPB is most closely related to *C. perfringens* δ-toxin (47% identity) and NetB (39% identity). It shares lower sequence homology to the staphylococcal toxins α-hemolysin (26% identity) and the bi-component leukocidins. Recently, we determined the molecular basis for the specificity of CPB towards endothelial cells by showing that PECAM-1/CD31, an endothelial and leukocytic adhesion molecule, serves as its cellular receptor (Bruggisser *et al*, 2020). So far however, no structural information has been available for CPB. In the present study, we describe the cryo-EM structure of CPB in SMA discs, which likely represents the membrane-inserted pore form, at near atomic resolution. We show that CPB forms a homo-oligomeric pore with a novel N-terminal β-barrel replacing the typical hemolysin latch domain. We propose that this β-barrel fold could be responsible for the efficient oligomerization of CPB in solution as well as influences the pore conductivity.

Our results have important implications for comparative studies on related toxins and the rational design of novel prophylactics and treatments directed against the action of clostridial hemolysin-like β-PFTs. In addition, the unique features of the newly discovered N-terminal β-barrel make CPB an attractive candidate for further studies into CPBs use in nanotechnology.

## Results

### Formation of the CPB oligomeric pore

To investigate the pore formed by CPB by single particle cryo-EM, we screened for a suitable detergent for oligomerization. CPB spontaneously assembles into SDS-resistant oligomeric species when stored in solution leading to partial precipitation of the protein (Supplementary Fig. 2a). Purification in presence of cholate followed by detergent removal and exchange with 2.5% SMA (Knowles *et al*, 2009) led to a homogeneous distribution of pores suitable for single particle cryo-EM (Supplementary Fig. 2b). Interestingly, the synthetic nanodiscs did not require addition of lipids suggesting that the SMA is able to directly wrap around the β-barrel of the CPB pore keeping the complex soluble. Particle distribution, orientation, and sample quality in SMA was better than either detergent or protein based nanodiscs (Supplementary Fig.3c). The average diameter of the particles was ∼100 Å. Two-dimensional classification of the particles revealed CPB pores with eight-fold symmetry. Further refinement and postprocessing resulted in an electron density map at an estimated 4 Å resolution (Figure 1a and Supplementary Fig. 3bd). This allowed to unambiguously build a model for the CPB octameric pore with the exception of two loops in the rim domain (Glu76 – Ser89, Ala283 – Pro287), 4 N-terminal amino acids and several amino acids with unresolved side chains (Figure 1b).

**Figure 1.**
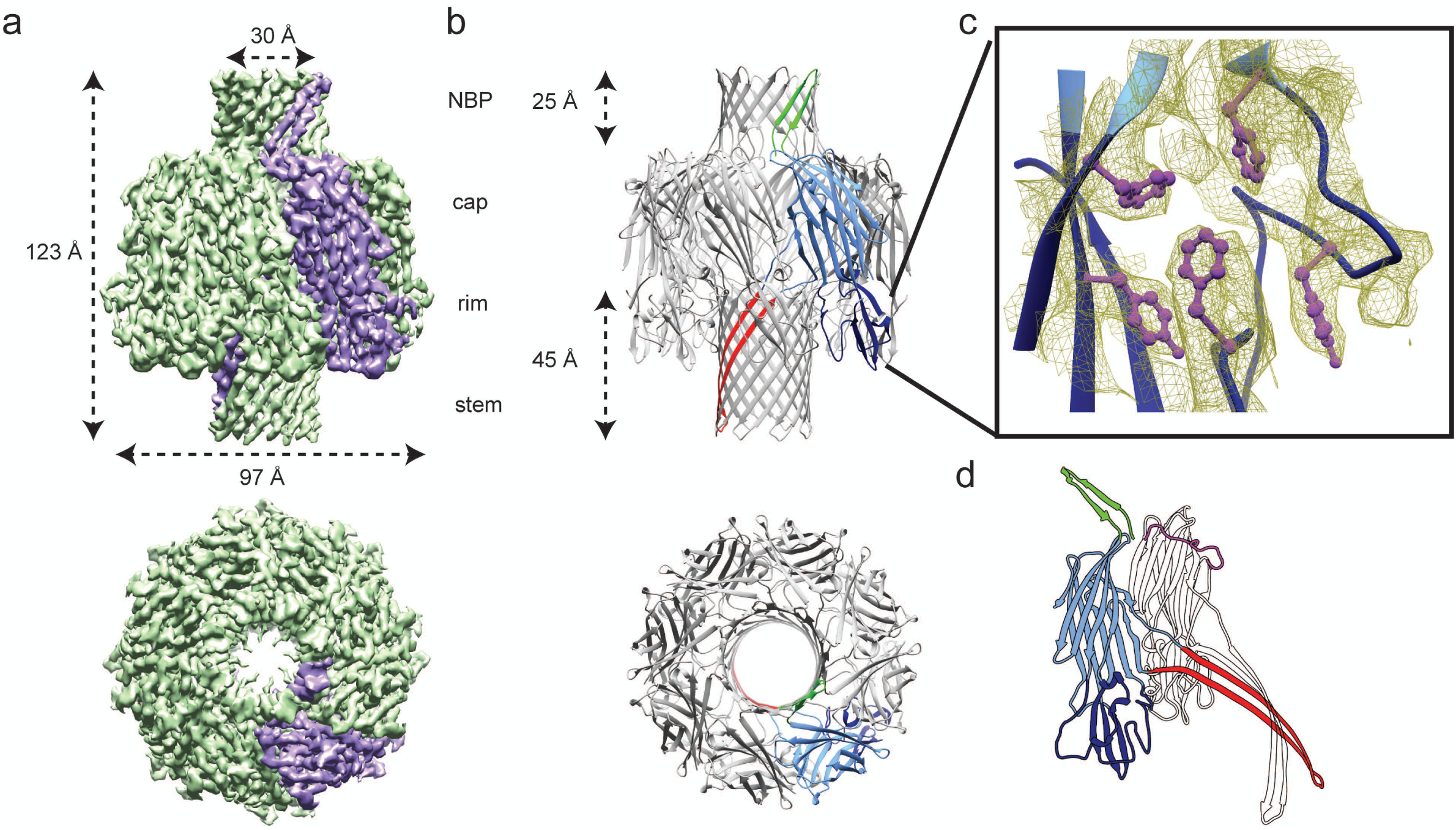
Structure of the CPB pore. (a) Side and top views of the 3.8Å sharpened cryo-EM map of the *C. Perfringens* β-toxin (CPB) with one monomer highlighted in purple. The map shows an extended 123 Å particle with two protrusions on both sides. The diameter of the particles is 97 Å with a visible channel of ∼30 Å diameter. (b) Model of the CPB oligomer showing a homo-octameric pore in top view and side view. The dimensions of the two β-barrels are indicated as measured from Cα to Cα. One protomer is colored by domains: green - N-terminal β-barrel protrusion; light blue - β-sandwich cap domain; dark blue - rim domain; red - stem domain. (c) Magnified view of the aromatic pocket of the rim domain docked in the cryo-em map showing the density of the aromatic side chains. (d) Cartoon model of the CPB protomer in the oligomer color coded as in (b) compared to a protomer extracted from the α-hemolysin oligomer. The N-terminus of α-hemolysin is highlighted in purple.

### Molecular Architecture of the CPB Pore

The cryo-EM analysis of CPB clearly showed that like other members of the α-hemolysin, the CPB protomer is composed of a cap, a rim, and a stem domain (Johnstone *et al*, 2021). In addition to the prototypical features of the family members, CPB possesses a second β-barrel on top of the cap domain. While the N-terminus of α-hemolysin is located inside the cap domain and wraps the vestibule-exposed surface of the adjacent protomer, the CPB N-terminus protrudes from the cap as a short hairpin, which assembles to form a 16-strand ß-barrel (Figure 1). We termed it the N-terminal β-barrel protrusion (NBP). The cap domain consists of a β-sandwich composed of two β-sheets and short α-helices. Two strands extend into the lower part of the molecule making up the rim domain. While the cap domain is one of the most conserved features within the α-hemolysin family, interesting differences are found in the rim domain, which is responsible for interactions with the membrane and protein receptors as seen by comparing two protomers extracted from the CPB and γ-hemolysin octamers (Supplementary Fig. 4). A homologous α-hemolysin phosphocholine binding pocket is present in the rim. However, at position 210, CPB is the only family member to have a bulky aromatic residue, a tyrosine, pointing inside the pocket (Figure 1c). Furthermore, CPB lacks a four-residue stretch, which lines the pocket. Among them a tryptophane, which is important for phosphocholine binding in staphylococcal hemolysins (Monma *et al*, 2004). These differences are compatible with the fact that the main receptor is a membrane protein rather than a lipid (Bruggisser *et al*., 2020). The loops at the base of the rim domain were well resolved except for two stretches of 14 and 5 residues, respectively (Glu76 – Ser89; Ala283 – Pro287). Both disordered stretches may contribute to a receptor binding surface (Supplementary Fig. 5). The stem domain is similar to the other family members. It contains a long, curved amphipathic hairpin, which is connected to the cap β-sandwich through two short coils forming the transmembrane β-barrel upon oligomerization.

The CPB octamer forms a 123 Å-long ring-like structure with a widest outer diameter of 97 Å (Figure 1). The channel runs along the 8-fold symmetry axis. It goes through the 25-Å long N-terminal ß-barrel, the central large 72.9 nm3 vestibule, formed by the cap domains, and the transmembrane β-barrel.

### N-terminal β-barrel protrusion

In order to further investigate the NBP domain and its role we mutated the first 23 amino acids of CPB and we constructed chimeras where the NBP was either deleted (Δ23CPB) or replaced by the equivalent N-termini of *S. aureus* α-hemolysin (Hla-Δ23CPB), γ-hemolysin component B (HlgB-Δ23CPB), or *C. perfringens* δ-toxin (δ-toxin-Δ23CPB). While the modification of the N-terminus seemed to partially affect protein expression and solubility in *E. coli* with a much lower protein yield for the chimeras when compared to the WT-CPB, the purified proteins still retained their ability to oligomerize and form pores (Supplementary Fig. 6). 2D classification of oligomers of the different mutants showed the lack of the NBP density for the Δ23CPB mutant as well as a lack of a structured N-terminus for the Hla-Δ23CPB and HlgB-Δ23CPB. The N-terminus from δ-toxin in the δ-toxin-Δ23CPB chimera seems to form an NBP similar to the WT-CPB suggesting that the N-terminus of δ-toxin oligomer might also adopt a β-barrel conformation in its oligomeric form (Figure 2a). Interestingly, the ability to form the NBP correlates with the cytotoxic activity as truncated Δ23CPB, HlgB-Δ23CPB, and Hla-Δ23CPB were inactive, whereas activity was fully restored for the δ-toxin-Δ23CPB (Figure 2b).

**Figure 2.**
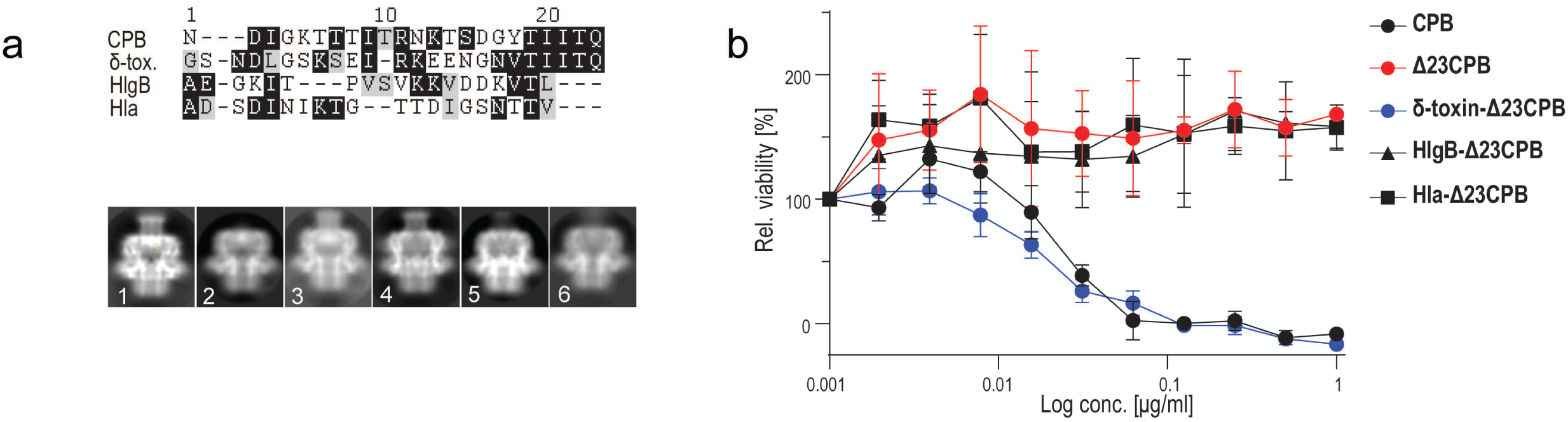
Investigating the N-terminal domain of different hemolysins. (a) Alignment of C. Perfringens β-toxin (CPB) N-terminus with N-termini of different hemolysins and cryo-EM 2D classification and average of N-terminally tagged WT CPB (1), Δ23CPB (2), δ-toxin N-terminus Δ23CPB (3), C-terminally tagged WT CPB (4), Hla N-terminus Δ23CPB (5) and HlgB N-terminus Δ23CPB (6). (b) Viability of HEK 293FT/CD31-GFP cells (transduced with CD31-GFP) as a percentage of untreated control cells after incubation with indicated concentrations of toxins (24 h, 37°C). Data are represented as means (n = 4) ± SD.

### Channel properties and dynamics

To investigate the mechanical and physical properties of CPB, we carried out atomistic Molecular Dynamics (MD) simulations of the whole toxin inserted into a lipid bilayer. The toxin’s channel features four constriction points in its two β-barrels (Figure 3a). The narrowest points have a ∼6 Å mean radius and are located within the NBP at the level of the positively charged Arg11 and Lys13. The channel reaches a maximum radius of about 20 Å within the vestibule of the cap. Quantifying the root mean square fluctuation (RMSF) of the toxin atoms revealed that the most flexible regions are the NBP and the intracellular mouth of the transmembrane β-barrel (Figure 3b). The NBP, while maintaining its overall structure, can oscillate off-axis, whereas the pore mouth can squeeze off a perfectly circular symmetry contributing to reducing the local pore radius (Supplementary Fig. 7). These observations are consistent with EM data obtained by performing a multibody analysis on the CPB particles and could explain the variability observed previously in the channel conductance (Shatursky *et al*, 2000) (Supplementary Fig. 3d). Examining charge distribution in the pore lumen revealed a high density of positive charges inside the β-barrel protrusion due to Arg11 and Lys13 (Figure 3). Potential mean force (PMF) profiles for Na+, Ca2+ and Cl- estimated via umbrella sampling indicated that this region should constitute an energy barrier for cations (4.6 kcal/mol for Na+). The wide cap vestibule features a balanced amount of positive and negative charges, while the transmembrane β-barrel is overall negatively charged, especially due to Glu152 and Glu162. These amino acids, associated with constriction points, determine a highly negative PMF for cations (as low as -24.6 kcal/mol for Ca2+) and a large 10.2 kcal/mol barrier for Cl- in agreement with previous reports suggesting that the CPB pore is mostly permeable to cations (Manich *et al*, 2008; Shatursky *et al*., 2000).

**Figure 3.**
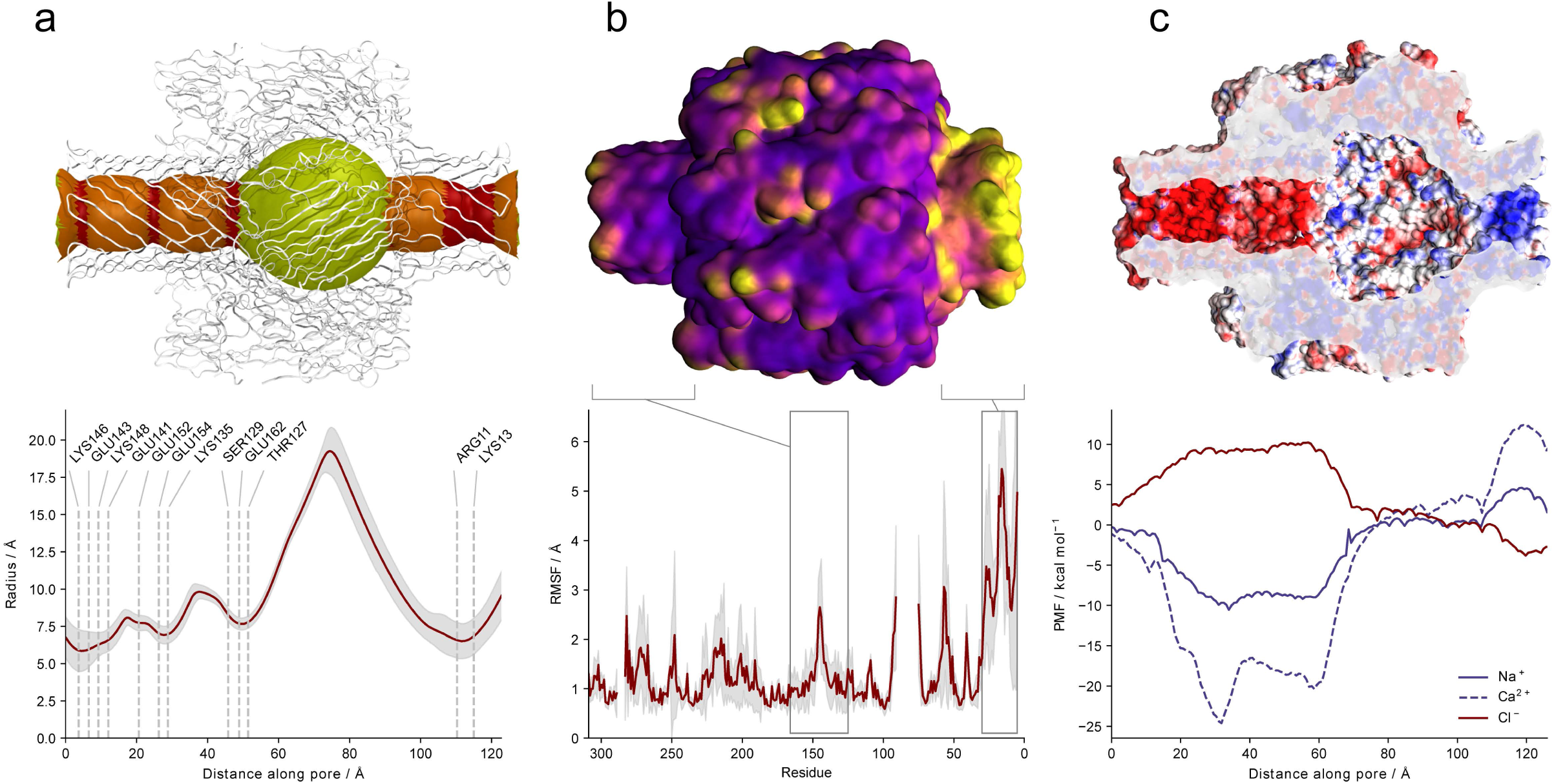
CPB dynamics and NBP measurements. (a) average pore radius in MD simulations. Above, regions with mean radius < 8 Å (constriction points) are shown in red, >8 Å and <10 Å in orange, and >10 Å in yellow. In the graph, mean values are shown in red, and standard deviations in grey. Polar amino acids at constriction points are annotated. (b) *C. Perfringens* β-toxin (CPB) root mean square fluctuation (RMSF) averaged over the eight CPB chains and three simulation repeats. Above, most mobile regions are shown in yellow, and least mobile in purple. In the graph, mean RMSF values are shown in red, and their standard deviation in grey. The N-terminal β-barrel Protrusion (NBP) is the most mobile region and, along with the flexible intracellular mount of transmembrane β-barrel, is annotated in the graph. (c) electrostatic properties of the pore internal cavity. Above, negative regions are shown in red and positive ones in blue. In the graph below, potential mean force profiles along the pore axis for Na+, Ca2+ and Cl- ions are shown. The NBP features a small positive region selective to anions, while the transmembrane β-barrel is expected to be highly selective to cations.

## Discussion

In this study we explored the structure of the membrane inserted oligomeric CPB and showed that it belongs to a new subclass of hemolysin-like β-PFTs with an additional β-barrel domain in its extracellular side. A detailed comparison of the N-terminal domains with structures of other oligomeric hemolysins is not possible. The N-terminus of the oligomeric γ-hemolysin is disordered (Yamashita *et al*, 2011). For NetB, oligomer crystallization was only possible after removing the first 20 amino acids of the protein (Savva *et al*, 2013). For *C. perfringens* δ-toxin only the monomer structure is available (Huyet *et al*, 2013) (Savva *et al*., 2013). In δ-toxin soluble monomer, unlike *S. auereus* α-toxin, the N-terminus adopts a β-hairpin conformation, which extends halfway along the cap β-sandwich and contacts the pre-stem (Supplementary Fig. 5a). Because of the high sequence similarity between CPB and *C. perfringens* δ-toxin (Supplementary Fig. 1 and Supplementary Table 1), it is reasonable to assume that the N-terminus of CPB adopts a similar conformation in the water-soluble monomer. To gain insight into the putative structure of CPB soluble monomer we made use of recent advances in protein structure prediction (Baek *et al*, 2021; Jumper *et al*, 2021). Both deep-learning methods generated predicted structures similar to the structure of δ-toxin monomer (Supplementary Fig. 5cd) with an N-terminus folded back and forming a β-hairpin. Comparing the AlphaFold predicted soluble monomer with a CPB protomer suggests putative rearrangements that may occur during oligomerization. In this model, the N-terminus initially forms an antiparallel β-sheet folded back against the first CPB β-strand of the cap domain likely stabilizing the soluble monomer. Following a local concentration increase and oligomerization, we hypothesize that the N-terminal β-hairpin swings by 90° and assembles in a β-barrel while the pre-stem domain refolds to form the transmembrane β-barrel (Figure 4 and Supplementary Movie 1). The net gain in hydrogen bonds resulting from the barrel formation is similar to α-hemolysin N-terminal latch and does not contribute significantly to the stability of the pore, which resists boiling in SDS with or without the NBP (Supplementary Fig. 6). Interestingly, in the predicted δ-toxin soluble monomer the N-terminus hairpin covers the subunit contact area at the cap level, which suggests that the CPB N-terminus may affect folding and oligomerisation (Supplementary Fig. 8). A similar role in regulating oligomerisation was also proposed for the N-terminal domains of the staphylococcal bi-component leukocidins and Hla (Jayasinghe *et al*, 2006; Yamashita *et al*., 2011).

**Figure 4.**
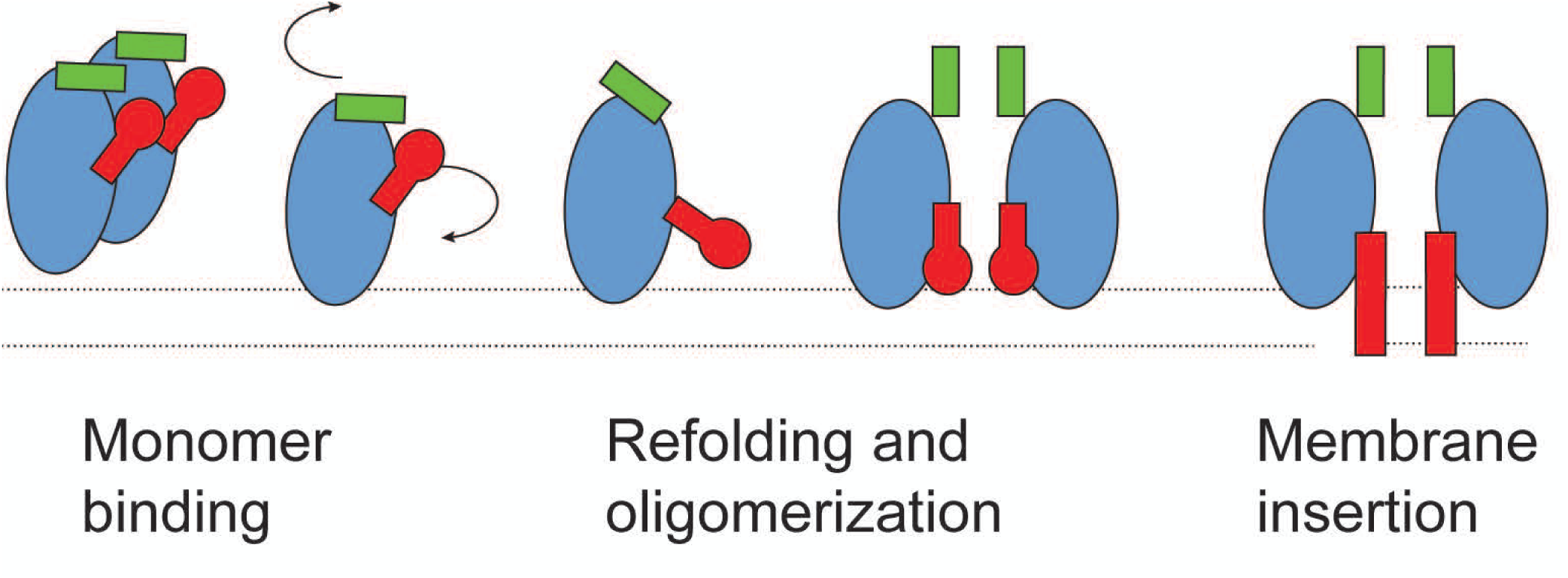
Putative model of CPB mode of action. Schematics of the predicted structural changes during *C. Perfringens* β-toxin (CPB) mode of action from the soluble inactive monomer to membrane-inserted oligomer subunits. Based on high level of sequence conservation with δ-toxin, we speculated that in soluble CPB the N-terminal hairpin (green rectangle) is located on top of the cap and rim domains (blue ellipsoid) in contact with the folded pre-stem domain (red club). We speculate further that the release of the pre-stem domain and membrane insertion coincides with the reorganization of the N-terminal hairpin, which through oligomerization becomes a N-terminal β-barrel.

In conclusion, the CPB pore has a C8 symmetry unlike the C7 symmetries of the homo-component members of the hemolysin family. It is the prototype of a new sub-group of the hemolysin-family characterized by a constriction at both ends of the channel forming a bipolar nanopore. The diameter of the two constrictions is smaller than for α-hemolysin (Song *et al*, 1996) and similar to the constriction present in the pore formed by MspA, a *M. Smegmatis* porin engineered for nanopore sequencing (Derrington *et al*, 2010). Our PMF profiles indicate that the NBP should constitute an energy barrier for cations while the transmembrane barrel has the opposite effect. The asymmetric, bipolar, nature of the CPB channel surface charge is a unique feature, which has not been observed to other hemolysin pores (Madai *et al*, 2019). This peculiar charge distribution could results in an increased flux of cations due to binding of anions to the NBP which could also explain the very low LD50 of CPB compared to other hemolysin pores. CPB bipolar charge distribution and unique pore architecture makes it an ideal candidate for small molecule sensing, while the double constriction connected by a large cavity could make it interesting for selective molecule delivery and transport. In addition, our deletions and N-terminus replacement suggests that the NBP selectivity can be easily altered without affecting the pore forming ability of the toxin.

Our findings about the structure of the CPB pore complex provides the basis for further studies on related PFTs as well as the structural basis of the interaction of β-PFTs with their respective receptor molecules. This could be exploited for the rational design of β-PFT inhibitors or anti β-PFT vaccines. The novel β-barrel protrusion observed here could also be present in other hemolysin-like family members, in particular from the clostridial proteins and elucidating its function could lead to a better understanding of this important toxin family.

## Materials and Methods

### Materials

Chemicals were purchased from Merck (Switzerland) or Sigma-Aldrich (Switzerland). Detergents were purchased from Anatrace (USA) or Sigma (Switzerland), SMA (SMALP 20010P and SMALP 50005P) from Orbiscope were a kind gift from the SMALP network and Polyscope, and oligonucleotides (Supplemental Table 2) were purchased from Microsynth (Switzerland).

### Molecular cloning

The codon optimized ORF (Genscript) encoding CPB was cloned as N-terminal His_6_ or N-terminal GST fusion into pET19-b (Novagen) using NcoI and BamHI sites. CPB containing a C-terminal His_6_-tag was generated using Q5® site-directed mutagenesis kit (NEB), pET-19b_His_6__CPB as template and primer pair #1 and elongation time of 4 min to delete the N-terminal His_6_, followed by a second PCR step using primer pair #2 and elongation time of 4 min to introduce the C-terminal His_6_. The His_6__CPB_Δ23_ construct was generated using Gibson Assembly® (NEB) and pET-19b_His_6__CPB was linearized using NcoI and BamHI sites. PCR product containing His_6_-CPB_Δ23_ was generated with primer pair #4, Ta and an elongation time of 1min. CPB chimera constructs were generated using Gibson Assembly® master mix (NEB) and pET-19b_His_6__CPB was linearized using NcoI and BamHI sites. PCR products containing the codon optimized N-terminal domains of δ-toxin (His_6_-CPB_Δ23_δ-toxin_(1-24)_), Hla (His_6_-CPB_Δ23_Hla_(1-20)_) and HlgB (His_6_-CPB_Δ23_HlgB_(1-19)_) were generated using pET-19b_His_6__CPB as template and indicated primer pairs, Ta and an elongation time of 1min (Supplementary Table 1). The resulting sequences were verified by Sanger Sequencing (Microsynth AG) and plasmids were transformed into *E. coli* Dh5α competent cells (NEB) for amplification.

### Recombinant toxin production and purification

The pET-19b plasmids encoding the CPB constructs were transformed into BL21 (DE3) strain (Sigma) for protein over-expression. Expression, solubility, and purification trials were done in small volumes of 0.05 - 0.2 l and upscaled to 3 – 9 l of LB medium (containing 100 μg/ml ampicillin and 1% glucose) according to the expression levels of the various CPB constructs. Expression cultures were inoculated with 1/50 volume of an overnight preculture grown from multiple colonies. Cultures were incubated at 37 °C and shaking at 180 rounds per minute. After growing to an OD_600_ of 0.6 – 0.8, expression cultures were cooled to 20°C and expression of CPB was induced by addition of 0.5 mM isopropyl-β-D-thiogalactopyranoside (IPTG). After protein expression (8 - 12 h at 20 °C), cells were pelleted 60 min at 3400 x g and 4 °C and stored at -20 °C until protein purification.

Cell pellet was resuspended in lysis buffer (50 mM Tris pH 8, 500 mM NaCl, 10mM imidazole, 1% TritonX-100) in the presence of lysozyme (0.2 mg/ml) and protease inhibitors (Sigma) followed by cycle high-pressure homogenisation (LM10 microfluidizer). Lysate was stirred on ice for 30 min in the presence of benzonase (Sigma) followed by two additional cycles of high-pressure homogenisation. After removal of cell debris by centrifugation for 60 min at 50’000xg (25’000 rpm using 45 Ti rotor) the supernatant was loaded on 5ml HiTrap chelating column (GE Healthcare, Germany) running on AKTA prime liquid chromatography system. The column was washed with 50 ml lysis buffer (containing 1% TritonX-100) and with 50 ml Buffer A (50 mM Tris, pH 8, 150 mM NaCl, 10 mM imidazole, 20-50 mM Na cholate). The protein was eluted with a linear gradient of imidazole (0 - 500 mM in 50 mM Tris pH 8, 500 mM NaCl, 20 mM Na cholate). Finally, fractions containing the oligomeric protein were dialyzed overnight (20 mM Tris pH 8, 150 mM NaCl, 20 mM Na cholate) and concentrated to 2 mg/ml.

For purification of monomeric CPB, the same *E. coli* strain was used but no detergent was added during the purification steps and a final concentration of 0.4 mg/ml was not exceeded because of protein precipitation due to spontaneous shift to oligomeric form at higher concentrations.

All purification steps were carried out at 4 °C. The purity of CPB preparations after each purification step was estimated by denaturing SDS-PAGE.

### Preparative size exclusion chromatography (SEC)

For further analysis by cryo-em preparative SEC was applied to separate monomeric CPB and other impurities from oligomers formed in detergent. The pooled and dialyzed protein solution from the metal chelate affinity chromatography was loaded on a 120 ml HiLoad® 16/600 Superdex® 75 pg column (GE Healthcare, Germany). The column was equilibrated in buffer (20 mM Tris, 150 mM NaCl, pH 8) run at a rate of 1 ml/min and eluted in 1 ml fractions. Fractions containing the CPB oligomer were pooled, concentrated and used for subsequent experiments.

### Sodium Dodecyl Sulphate Polyacrylamide Gel Electrophoresis (SDS-PAGE)

Samples were mixed with 1/5 of their volume of 5x SDS sample buffer and incubated at RT for at least 5 min (or 5 min at 95°C if referred to as boiled) prior to loading on a gel. 10 μl of samples were loaded on 10% or 16% polyacrylamide gels and subjected to electrophoresis at limiting current of 55 mA for 45 min. Gels were stained in a 0.2% Coomassie Brilliant Blue G solution for 20 min or fixed in prefixing solution and stained with the sensitive colloidal staining solution.

### Blue Native Polyacrylamide Gel Electrophoresis (BN-PAGE)

The samples (10 μg protein/lane) after a clarifying spin (20,000 x g, 15 min, 4 °C), were mixed with 5x BN sample buffer (2.5% (w/v) Coomassie brilliant blue G-250, 100 mM Bis-Tris, 250 mM 6-aminocaproic acid, 50% Glyerol, pH 7.0) and analysed by electrophoresis in a blue native gel containing 4 - 16% gradient of acrylamide (29:1 Acrylamide:Bis) using the SE 600 Vertical electrophoresis system (GE Healthcare Life Sciences). The gels were run at 4 °C at constant 200 V for 2 h followed by 6 h at 600 V with max. 20 mA. The mixture NativeMark Unstained Protein Standard (Thermofischer) was used to monitor the migration of molecular weight marker proteins. Gels were stained with colloidal Coomassie.

### Negative staining

400 mesh carbon coated copper grids (CF400-Cu Electron Microscopy Sciences) were glow discharged using a CCU-010 sputter/carbon coater (Safematic) with negative polarity (10 mA) for 45 s immediately before usage. 4 µl of the protein sample was adsorbed to the prepared grids for 45 s. After drying excess liquid with a filter paper (blotting), the samples were washed six times with ddH2O and stained immediately by placing the grid on top of a drop of 2% uranyl acetate solution for 1 min. Finally, the stained grid was blotted dry and dried completely under a hood prior analysis. Electron micrographs were recorded at 105’000 x magnification by an Olympus-SIS Veleta CCD camera using a Tecnai Spirit G2 electron microscope (FEI, USA) operating at an acceleration voltage of 80 kV.

### Sample preparation for cryo-EM

SMAs are synthetic copolymers composed of styrene (S) and maleic acid anhydride (MA) that function as an alternative to detergents and amphipols (Dorr *et al*, 2016). They can be used to directly purify a membrane protein from their natural lipid environment forming SMA lipid particles (SMALPs) in which the membrane proteins are surrounded by a small disk of lipid bilayer encircled by polymer, similarly to MSP nanodiscs. We tested the SMA (S:MA ratio 2.3:1 and 1.4:1) directly as substitute for detergents Big-Chaps and sodium cholate, without the addition of lipids. Therefore, SMA at 2.5% (w/v) was incubated with oligomeric CPB at a concentration of around 1.5 mg/ml for 1h at RT and detergent was removed with Amberlite® XAD®-2 biobeads (Sigma) over night at RT. The formed SMA CPB particles were concentrated to around 4 mg/ml and analysed by single-particle cryo-EM following gel filtration.

### Specimen preparation and data collection

For cryo-EM 3ul of the protein sample at different concentrations were deposited onto a copper grid (quantifoil Cu 200 mesh R2/1, R1.2/1.3) that was glow discharged 10-20” 10mA using a Baltzers CTA 010. Vitrification was performed by plunging into liquid ethane in an atmosphere at 4°C and 100% humidity using a Vitrobot Mark IV. Vitrified grids were stored in liquid nitrogen prior to acquisition which was performed on a FEI Tecnai F20 equipped with a Falcon III camera. Images were recorded as stack of frames using FEI EPU automatic data collection with a total dose not exceeding 60e^-^/Å^2^ and processing was performed in RELION (Zivanov *et al*, 2018). Acquisition was performed over several days using an in-house liquid nitrogen filling robot for the side entry holder (Gatan 626) which was set to refill the Dewar every 3h. Post-acquisition the recorded movie frames were motion corrected and summed using Motioncor followed by CTF correction using Ctffind (Rohou & Grigorieff, 2015; Zheng *et al*, 2017). A small subset (less than 2000 particles) was selected by hand and an initial 2D classification was used to generate the references used for autopicking 1874390 particles. Autopicking was followed by 2D classification of 4x binned particles without image alignment which resulted in an initial subset of 792433 particles. A new round of 2D classification with image alignment on the initial subset resulted in 382962 particles which were re-extracted unbinned. 2D classification of the unbinned data set allowed us to select the best particles suitable for high resolution. The best data set selected contained 260481 which could be refined to an estimated ∼4.2 Å resolution as estimated by RELION. Performing the implemented Bayesian polishing step led to an improved resolution of 3.8 Å.

### Model building and refinement

Model building inside the electron density map was based on HlgAB (PDB: 3b07) (Yamashita *et al*., 2011) and was performed in Coot and Phenix using real space refine (Table 1) (Emsley *et al*, 2010; Liebschner *et al*, 2019). Several models were generated for visualization purposes, the CPB octamer model with outside chains for residues lacking a clear EM density in the map and a complete model where the residues with missing EM density were fitted with the most appropriate rotamer and then inspected in coot and refined to remove clashes. A third model was generated fitting the missing loops using Phenix implementation for fit loops (Supplementary Fig. 5b). All images were generated using Chimera, Coot and PyMol (Emsley *et al*., 2010; Pettersen *et al*, 2004; Schrodinger, 2015).

**Table 1.** Refinement statistics. **Cryo-EM data collection, refinement and validation statistics**

|  |  |
| --- | --- |
|  | #1 CPB<br>D_1292117811<br>(EMDB-xxxx)<br>(PDB xxxx) |
| **Data collection and processing** |  |
| Magnification | 80000 |
| Voltage (kV) | 200 |
| Electron exposure (e-/Å<sup>2</sup>) | 60 |
| Defocus range (μm) | -1.8 – -3.0 |
| Pixel size (Å) | 1.306 |
| Symmetry imposed | C8 |
| Initial particle images (no.) | 382962 |
| Final particle images (no.) | 260481 |
| Map resolution (Å) | 3.8 |
| FSC threshold | 0.143 |
| Map resolution range (Å) | 3.6 – 7.5 |
| **Refinement** |  |
| Initial model used (PDB code) | 3B07 |
| Model resolution (Å) | 4/0.143 |
| FSC threshold |  |
| Model resolution range (Å) | 3.8/4.5 |
| Map sharpening *B* factor (Å<sup>2</sup>) | -266 |
| Model composition | 15840/2232/0 |
| Non-hydrogen atoms |  |
| Protein residues |  |
| Ligands |  |
| *B* factors (Å<sup>2</sup>) | 266 |
| Protein |  |
| Ligand |  |
| R.m.s. deviations | 0.36 Å/0.54° |
| Bond lengths (Å) |  |
| Bond angles (°) |  |
| Validation | 4/0/0 |
| MolProbity score |  |
| Clashscore |  |
| Poor rotamers (%) |  |
| Ramachandran plot | 97/3/0 |
| Favored (%) |  |
| Allowed (%) |  |
| Disallowed (%) |  |

### Molecular Dynamics

Three independent molecular dynamics simulation repeats were performed by embedding the atomistic CPB octamer into a POPC bilayer, solvating it with TIP3P water and charge-balancing the resulting system with Na+ counterions. All simulations were parameterized using the Amber14SB (Maier *et al*, 2015) forcefield with Slipids lipid parameters (Jambeck & Lyubartsev, 2012), and performed with the GROMACS engine (Abraham *et al*, 2015). To equilibrate the systems, the temperature was brought up to 310 K over 1 ns with the V-rescale themostat (NVT conditions). The pressure was then set to 1 bar over 1 ns (NPT conditions) using the Berendesen barostat. During these first equilibration phases, we promoted the relaxation of the flexible NBP region into a conformation of minimal energy by restraining the distances between atoms expected to form an inter-chain H-bond. These restraints were then slowly lifted over 15 ns NPT simulation using the Nose-Hoover thermostat and Parinnello-Rahman barostat. Finally, 200 ns free NPT simulations were carried out, and conformations collected every 0.1 ns. During all simulations, electrostatics were calculated with the particle mesh Ewald algorithm, and the LINCS algorithm was used to enable the use of a 2 fs time step. Assessment of internal pore diameter calculations were performed with HOLE2 (Smart *et al*, 1996) on each simulation conformation. The root mean squared fluctuation of each atom was calculated over an aggregation of all independent trajectories, and then cast onto a volume representation to produce the visualisation of Figure 3. Potential mean force (PMF) profiles along the pore z-axis for Na+, Ca2+ and Cl- ions were calculated via umbrella sampling. To this end, ions were restrained with a 1000 kJ nm-2 harmonic potential along the axis with a 2 Å spacing, and simulated at each position for 10 ns. We then reconstructed the one-dimensional PMF profile along the pore using the weighted histogram analysis method (WHAM). The electrostatics of the internal pore cavity were visualised by casting the continuum electrostatic potential calculated with the adaptive Poisson-Boltzmann solver (APBS) (Jurrus *et al*, 2018) onto a simulated volumetric map of CPB. Volumes simulations and image rendering were produced with VMD (Humphrey *et al*, 1996).

### Cytotoxicity Assay

The HEK 293FT/CD31-GFP cells were previously established by lentiviral transduction of HEK 293FT cells with lentiviral plasmid encoding mouse CD31-GFP (Bruggisser *et al*., 2020) rendering them sensitive for CPB. HEK293FT/CD31-GFP cell lines were cultured in DMEM medium (Gibco, product 41965-039) supplemented with 10% fetal calf serum FCS (Gibco), 10 mM Hepes pH 7.2 (Gibco), 4 mM L-Glutamine (Gibco), in the presence of penicillin-streptomycin (Gibco) and puromycin (1 mg/ml), grown at 37°C in an atmosphere containing 5% CO2.

### Effects of CPB and CPB mutants on cells were measured by using Resazurin assay

Cells (2 × 104 cells/ml) grown to confluency in a 96 well plate were incubated with CPB (1 μg/ml CPB starting concentration, 1:2 dilution steps) for 24 h. Resazurin dye was added to a 0.002% final concentration, incubated for 4 h at 37°C and fluorescent signal intensity was quantified using the EnSpire Multimode Plate Reader (PerkinElmer) at excitation and emission wavelengths of 540 and 612 nm respectively.

## Acknowledgements

We wish to thank Marek Kaminek for indispensable assistance with all the EM equipment. JB wishes to acknowledge Michael Stoffel and Véronique Gaschen for their help with negative stain preparations. We wish to thank Durham HPC Hamilton for computational resources. Images were acquired on an instrument supported by the microscopy imaging center (MIC) of the University of Bern. Funding: This study was supported by a grant of the Berne University Research Foundation to II, by the Swiss national research foundation grant 31003A_179520 to BZ, by the Swiss national research foundation grant 31003A_169381 to HP, and by the Engineering and Physical Sciences Research Council fellowship EP/P016499/1 to MTD.

## Author contribution

JB and II equally contributed to this study. JB did cloning and performed detergent screening. JB and II performed sample preparation. II performed cryo-EM acquisition, image processing, and model building. SCM carried out molecular dynamics simulations. JB, II, and BZ, SCM and MTD analysed the data. JB, and II, SCM and MTD prepared the figures. JB and II wrote the initial manuscript draft.

All authors contributed to manuscript writing. MTD, BZ and HP supervised the study and acquired funding.

## Data availability

The EM map and the pdb model of the CPB oligomer are available as PDB ID 7Q9Y and EMD-13876.

## ABBREVIATIONS

CPB: C. perfringens β-toxin
SMA: styrene maleic acid
PFT: pore forming toxin
NBP: N-terminal β-barrel protrusion
PMF: potential mean force
RMSF: root mean square fluctuation

---

**Supplementary Table 1.**
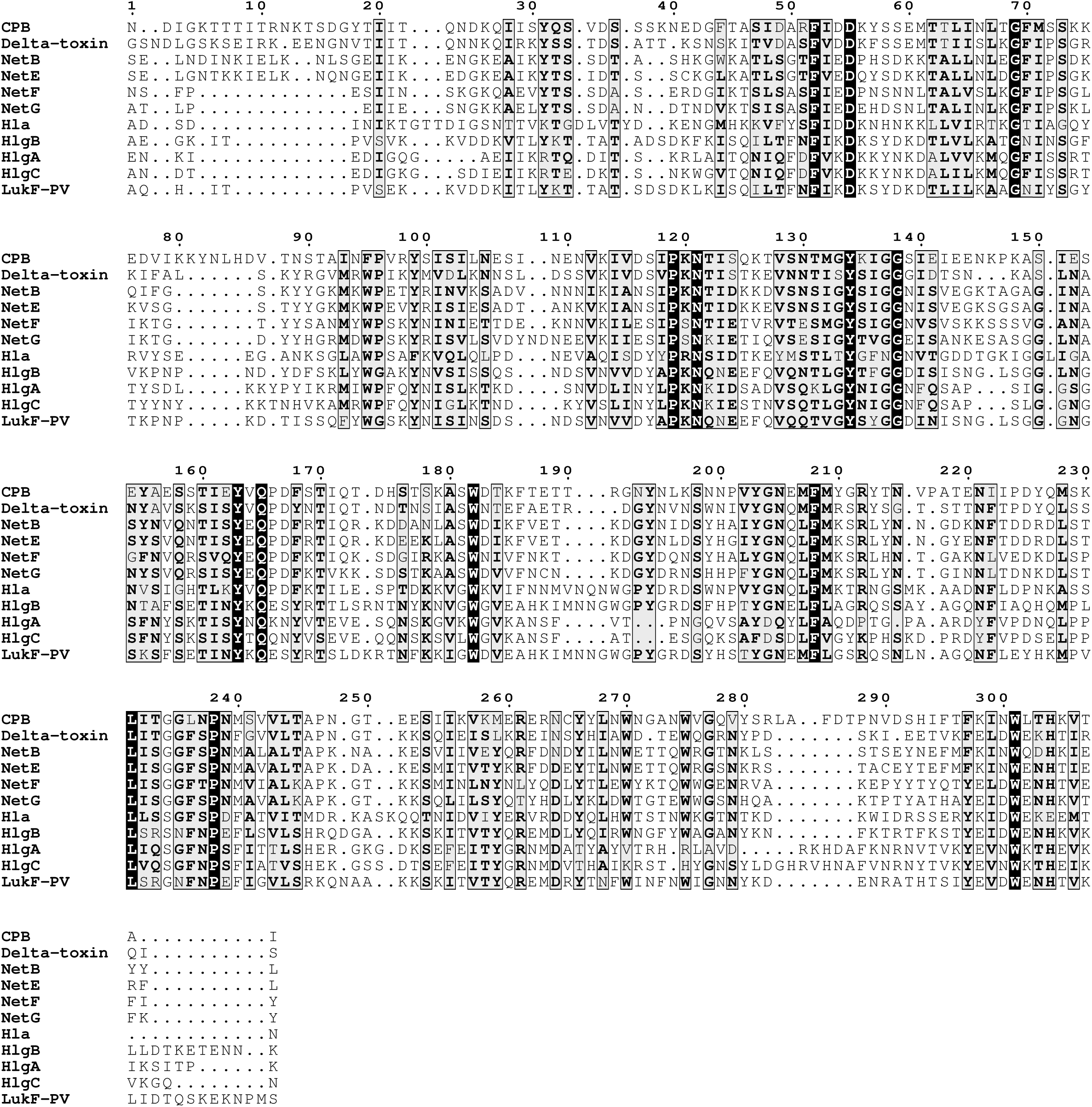
Sequence Alignment Sequence alignment of CPB and related toxins. CPB (Uniprot ID: Q9L403), Delta toxin (Uniprot ID: B8QGZ7), NetB (Uniprot ID: A8ULG6), NetE (Uniprot ID: A0A0D3QGW9), NetF (Uniprot ID: A0A0D3QGV4), NetG (Uniprot ID: A0A0D3QH83), and representative sequences from the S. aureus, including leukocidin S components: HlgA (Uniprot ID: P0A074), HlgC (Uniprot ID: Q07227), F components: LukF (Uniprot ID: Q5FBD2), HlgB (Uniprot ID: P0A077) and hemolysin Hla (Uniprot ID: P09616). Multiple sequence alignments were done using T-coffee and figure has been made using ESPript program (DOI: 10.1006/jmbi.2000.4042; DOI: 10.1093/nar/gku316).

**Supplemental Table 2.**
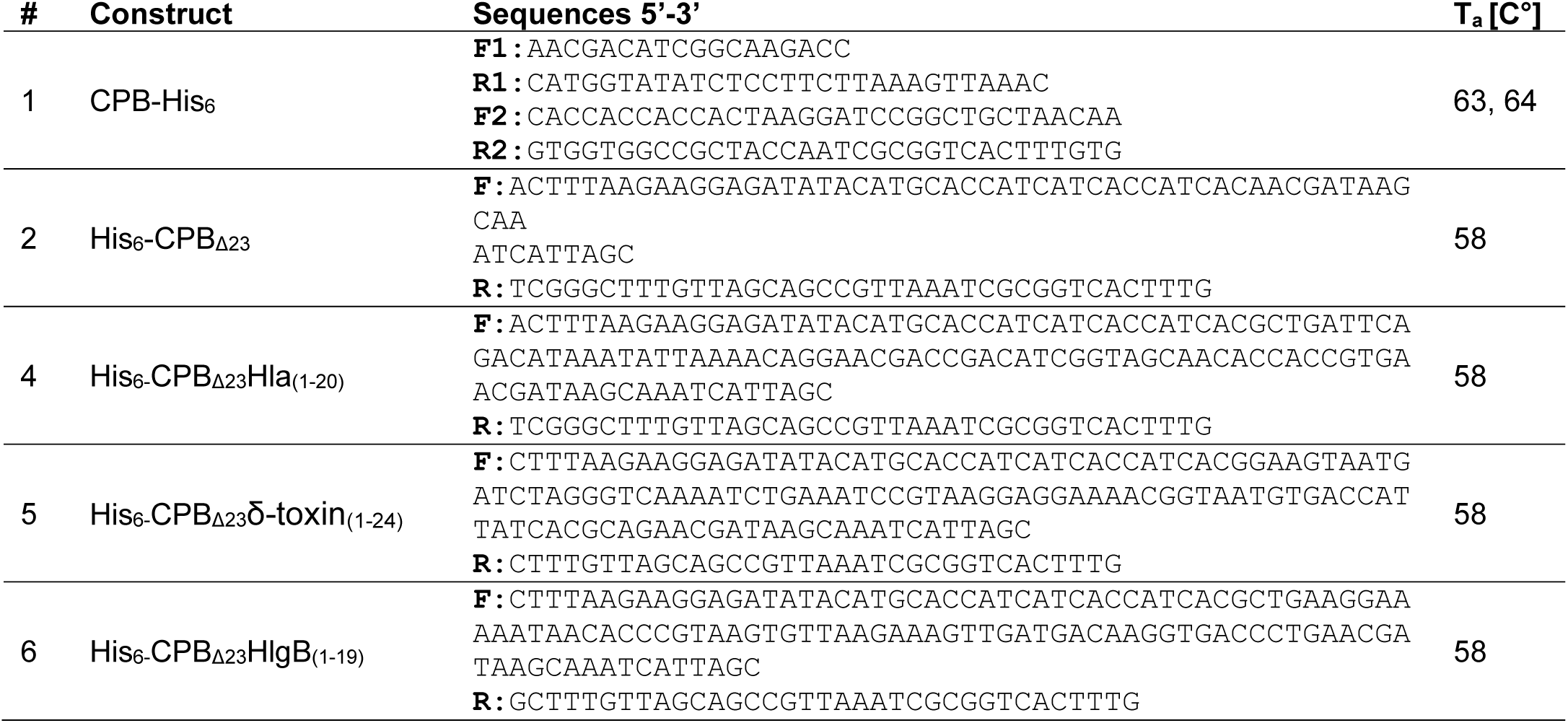

**Supplemental Figure 1.**
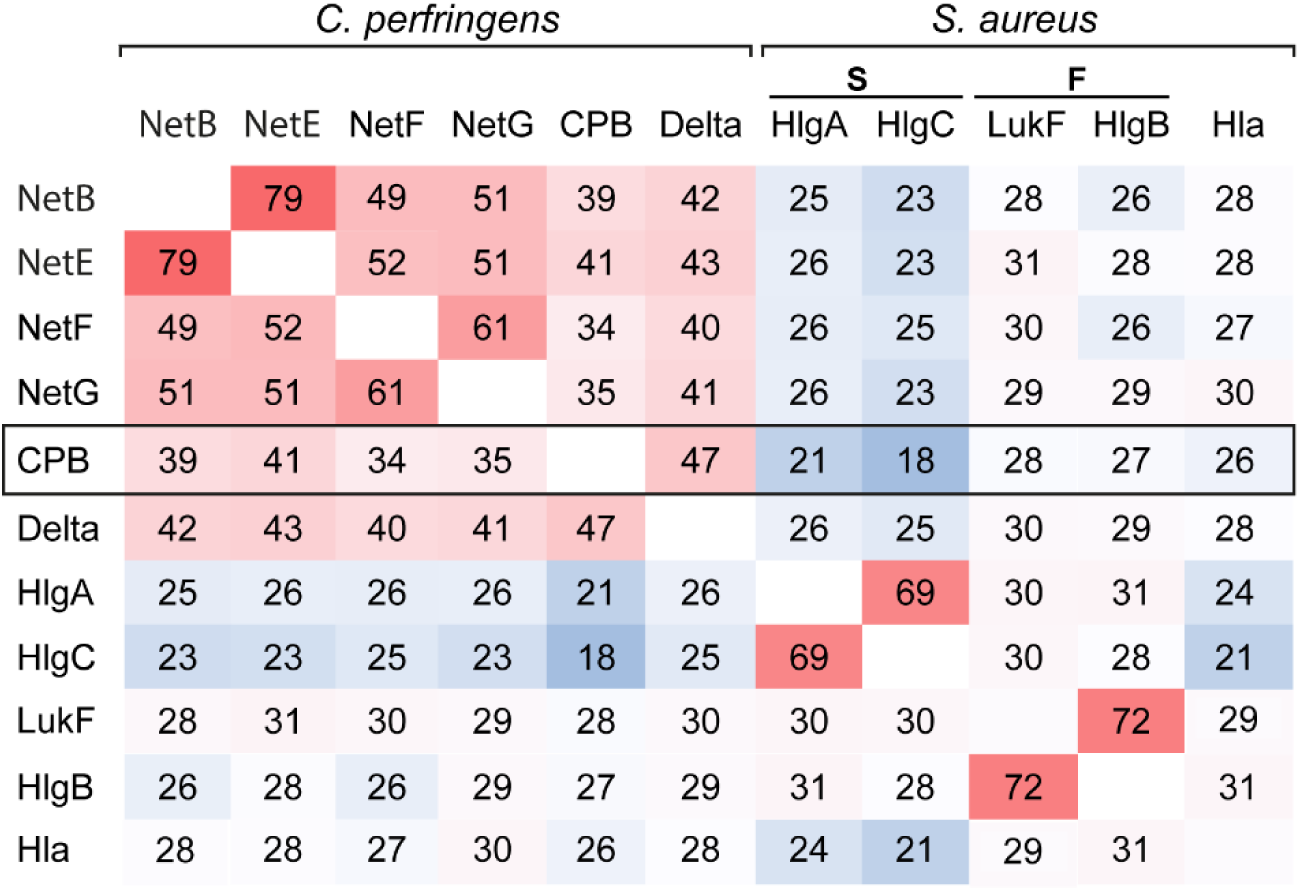
Sequence homology between the different hemolysin family members showing CPB highest similarity to the clostridial δ-toxin. Heatmap shows percent identity matrix of protein alignments, colors correspond to the percent identity with high values (red), medium values (white) and low values (blue). The Identity matrix was done using Clustal Omega (Madeira et al., 2019). NetB (Uniprot ID: A8ULG6), NetE (Uniprot ID:A0A0D3QGW9), NetF (Uniprot ID: A0A0D3QGV4), NetG (Uniprot ID: A0A0D3QH83), CPB (Uniprot ID: Q9L403), Delta toxin (Uniprot ID:B8QGZ7) and representative sequences from the S. aureus, including leukocidin S components: HlgA (Uniprot ID: P0A074), HlgC (Uniprot ID: Q07227), F components: LukF (Uniprot ID: Q5FBD2), HlgB (Uniprot ID: P0A077) and hemolysin Hla (Uniprot ID: P09616).

**Supplemental Figure 2.**
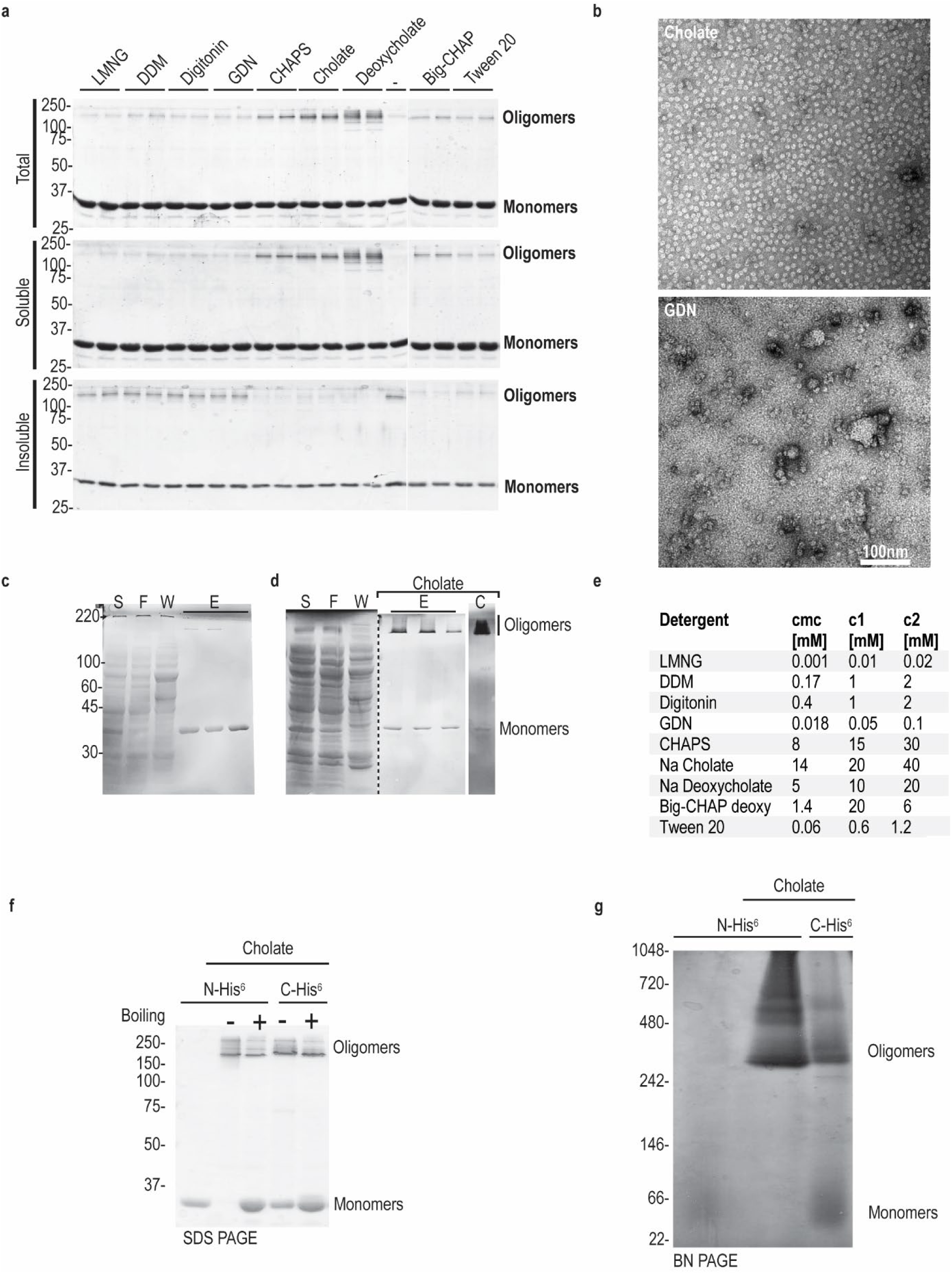
Oligomerization and solubility of CPB oligomers in different detergents showing cholate and deoxycholate as the best candidates for purification and solubilization of CPB pores (a). Each detergent was added to the CPB sample for 16h at 4° and then the sample was centrifuged at 15k xg for 10 minutes. The pelleted insoluble fraction was recovered in sample buffer and loaded on SDS-PAGE together with the soluble fraction and the initial (total) fractions. Negative stain (b) of representative CPB oligomers after purification in 30mM cholate with evenly spread oligomers (top) vs after exchanging cholate with 0.1mM GDN with aggregates and background particles (bottom). SDS-PAGE gels of His6-CPB purification in the absence (c) and presence (d) of cholate. 10μl of supernatant (S), flow through (F) and elution (E) fractions were loaded and gels stained with coomassie stain. Most oligomers precipitate in the absence of detergent during the purification, whereas after concentration (C) almost all CPB shifted into the oligomeric state in the presence of cholate. Table of the detergents used for the screen, their CMC and their concentration (e). Characterization of N- and C-terminal tagged CPB constructs by gel electrophoretic analysis under denaturing (f) and native (g) conditions. SDS-PAGE gel analyis of CPB samples (1μg per lane) showing monomeric CPB in the absence of detergent and SDS resistant oligomers in the presence of cholate. CPB samples containing cholate were boiled for 5 min at 95°C (+) or not (-).

**Supplemental figure 3.**
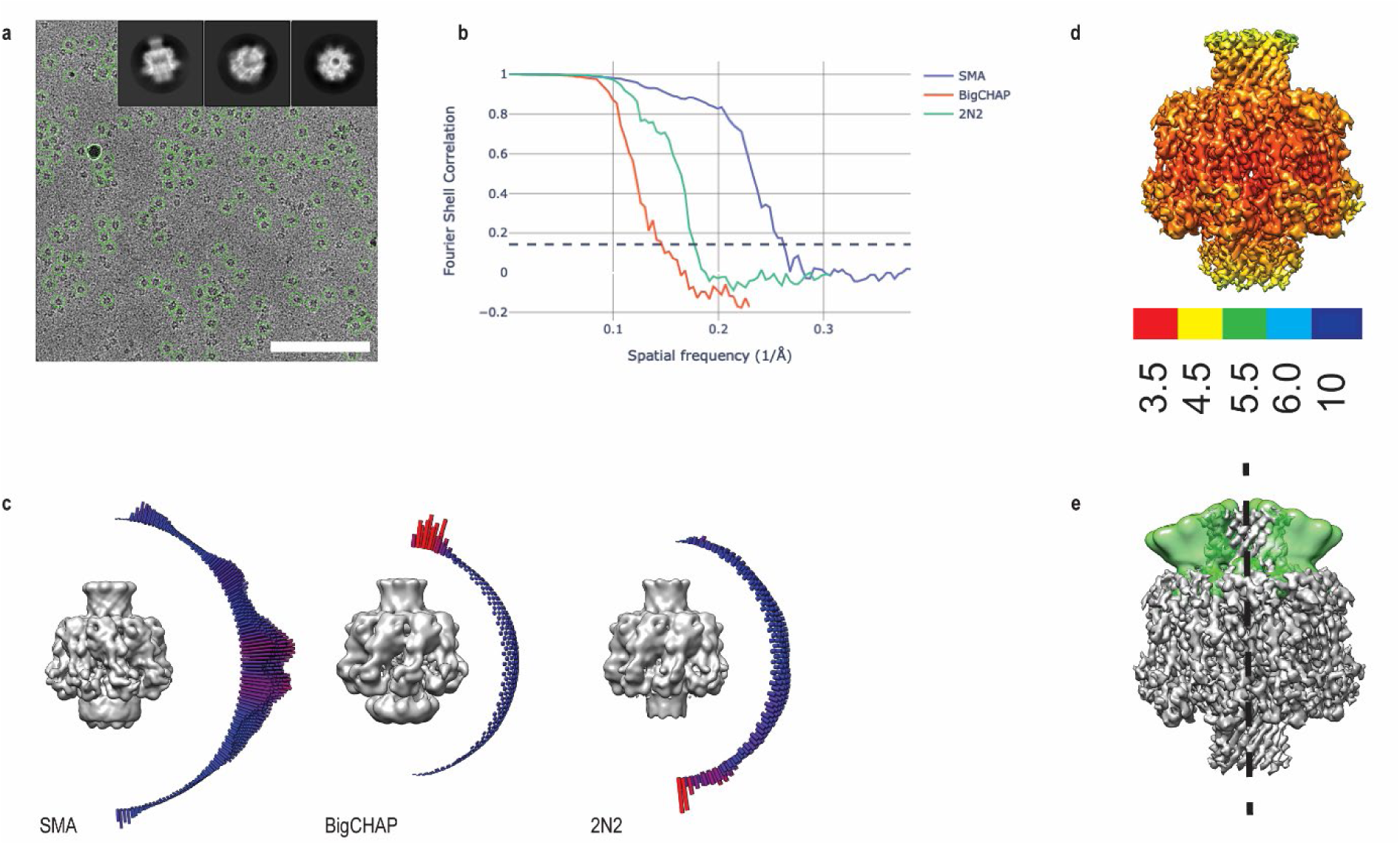
a) Electron micrograph of a typical field of view of oligomeric *C. Perfringens* β-toxin (CPB). Automatically picked particles shown in green. Inset showing characteristic 2D class averages, side, tilted, top. Scale bar is 100 nm. b) FSC showing the resolution of CPB oligomers in different reconstruction conditions with SMA giving the best orientation of the sample on the grid c) Refined cryo-EM maps and angular distribution of particles of CPB reconstituted in SMA, BigCHAP or 2N2 nanodiscs. (d) Local resolution estimate performed in relion showing the cap and rim domains at the highest resolution. Position of the NBP at the extremes of the multibody analysis (e) performed in relion (in green) compared to the CPB map(gray). (d) Putative rearrangements required to convert from a soluble CPB monomer to oligomer. NBP is shown in green, cap domain in cyan, rim domain in dark blue and the stem in red.

**Supplemental figure 4.**
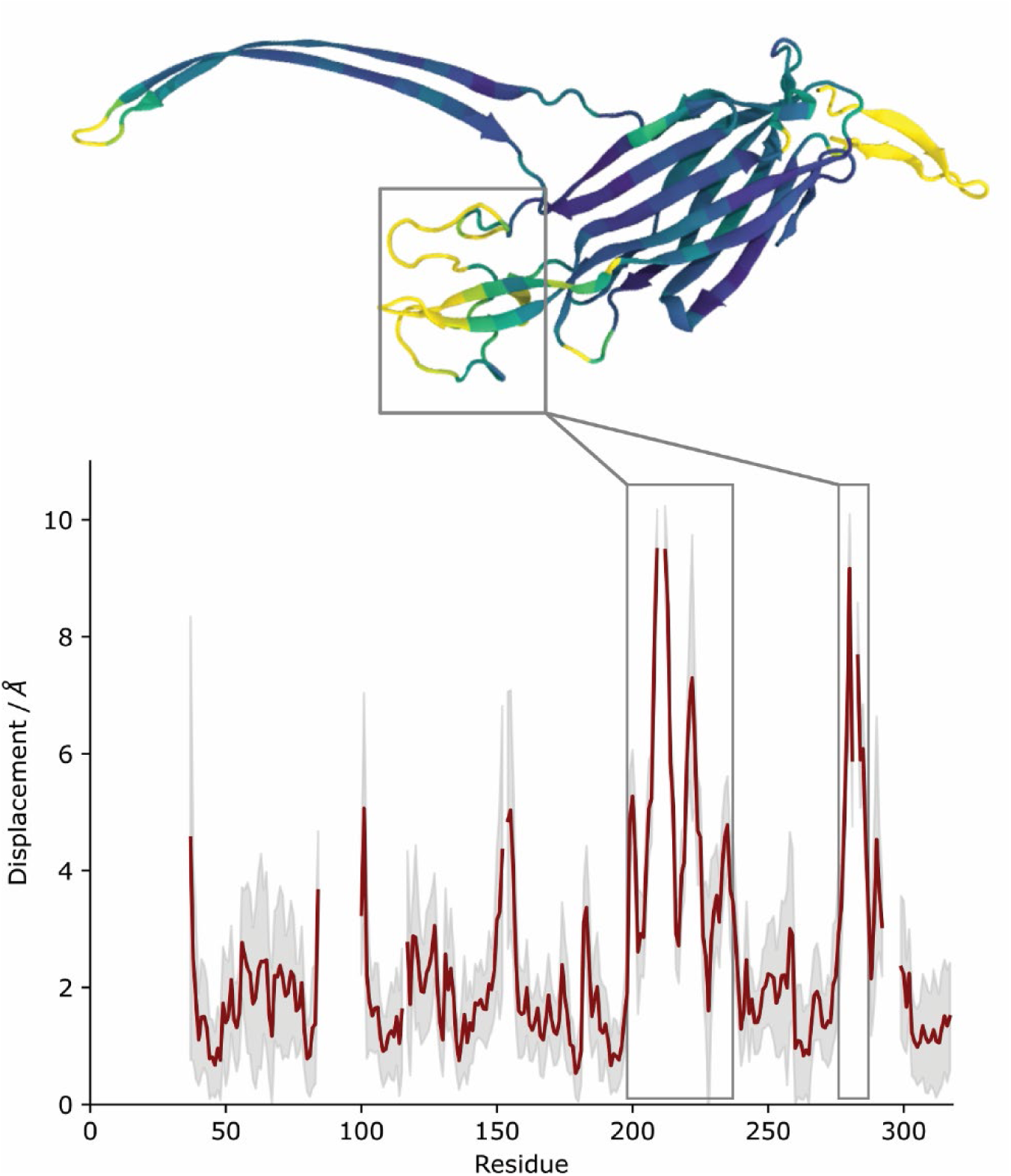
Monomers extracted from the *C. Perfringens* β-toxin (CPB) and γ-hemolysin (PDB: 3B07) octamers were aligned (STAMP structural alignment), and the Euclidean distance of their paired Cα atoms measured. In the graph, the black line reports on the distances measured when using our CPB structure. The red line reports on the mean difference when using all structures extracted from our MD simulations for comparison, and the gray shaded region reports on the standard deviation. The protein rendering is coloured according to the mean difference of simulated structures.

**Supplemental Figure 5.**
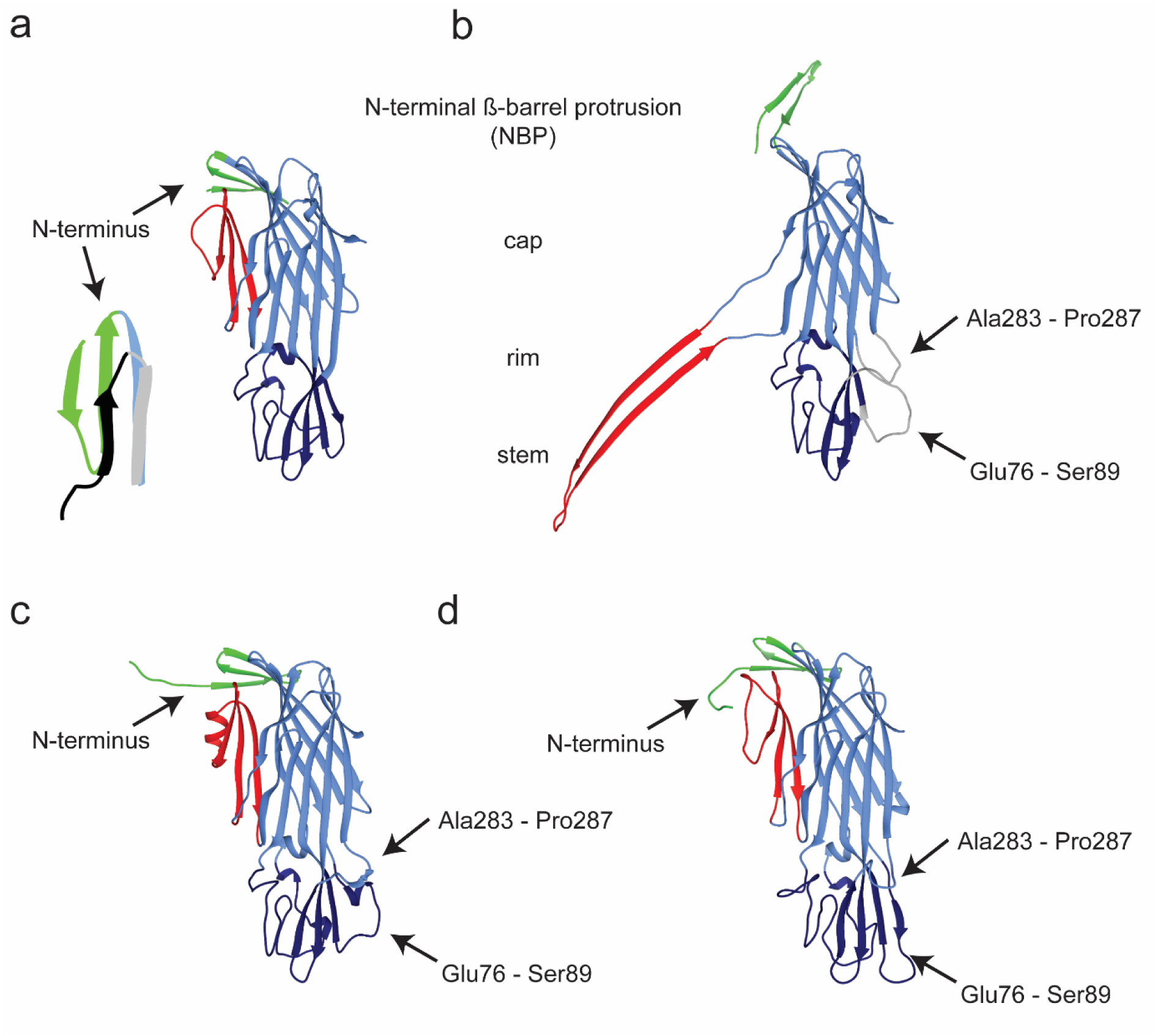
Model of δ-toxin (2YGT) using the same color code as in Figure 1 showing the position of its N-terminus folded back as an additional strand (a) with the inset showing a comparison between the N-terminus of the δ-toxin (green) and the N-terminus of α-toxin (black – 4YHD). Model of the monomeric CPB extracted from the oligomer structure color coded as before (b). The missing loops in the cap and rim domain are modeled and shown in gray. The missing loops are only shown as visual guide for the number of missing amino acids in the model as the map quality in those regions is not good enough for model building. Prediction of the CPB monomer structure by AlphaFold (c) and RosettaFold (d) color coded as before. The predicted N-terminus folds as the Nterminus as δ-toxin. The missing loops in the rim domains fold similarly to our modelling (b) in the case of alphafold algoritm while RosettaFold modelling extends the longer missing loop to a long β-sheet.

**Supplemental Figure 6.**
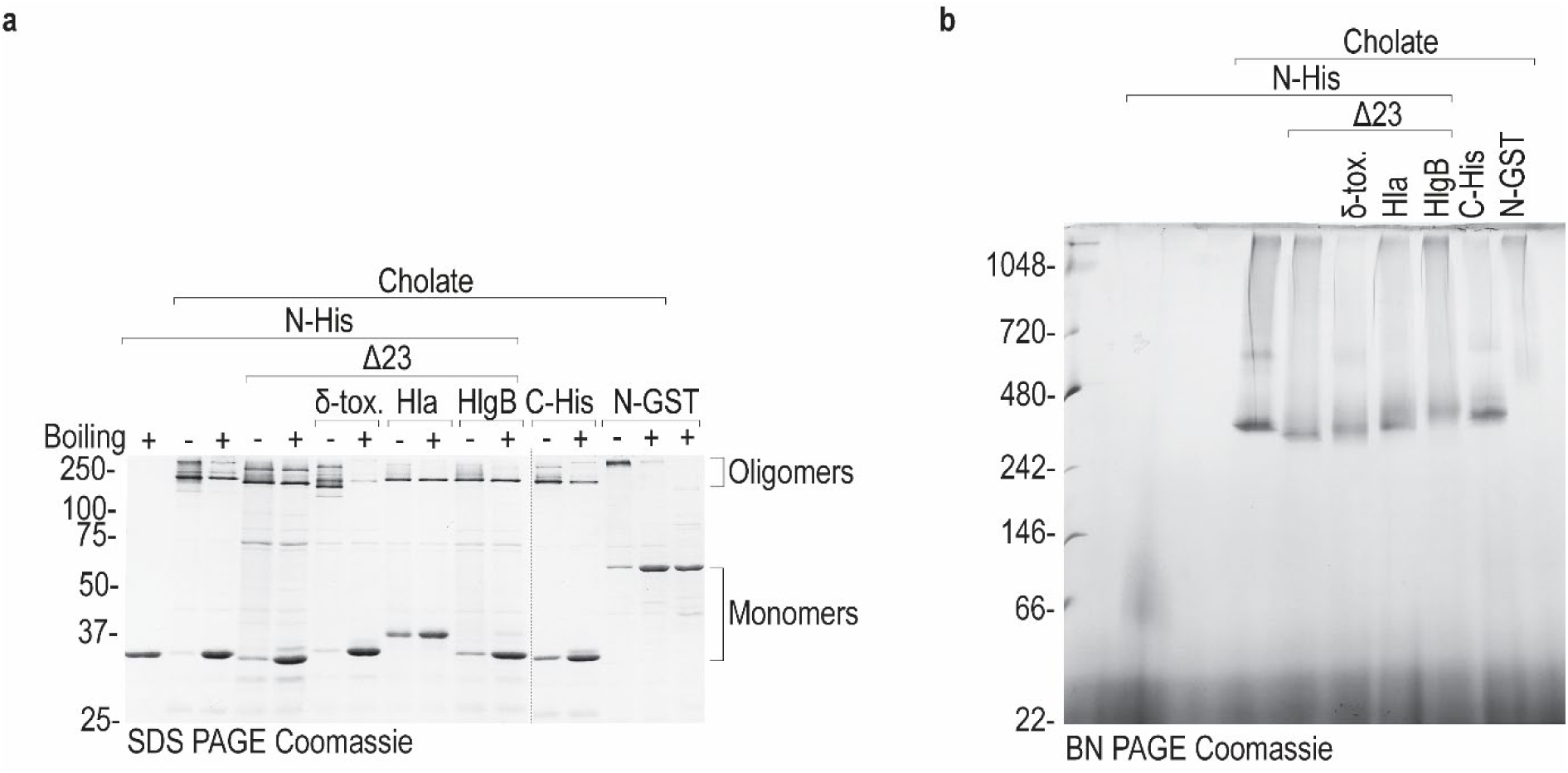
Characterization of CPB constructs with different N-termini by gel electrophoretic analysis under denaturing (a) and native (b) conditions. SDS-PAGE gel analyis of CPB samples in cholate (1μg per lane). CPB samples containing cholate were boiled for 5 min at 95°C (+) or not (-). (b) Coomassie stained Blue native PAGE gel (4-16%) of CPB samples (10ug per lane) showing oligomer formation for all different N-termini constructs.

**Supplemental Figure 7.**
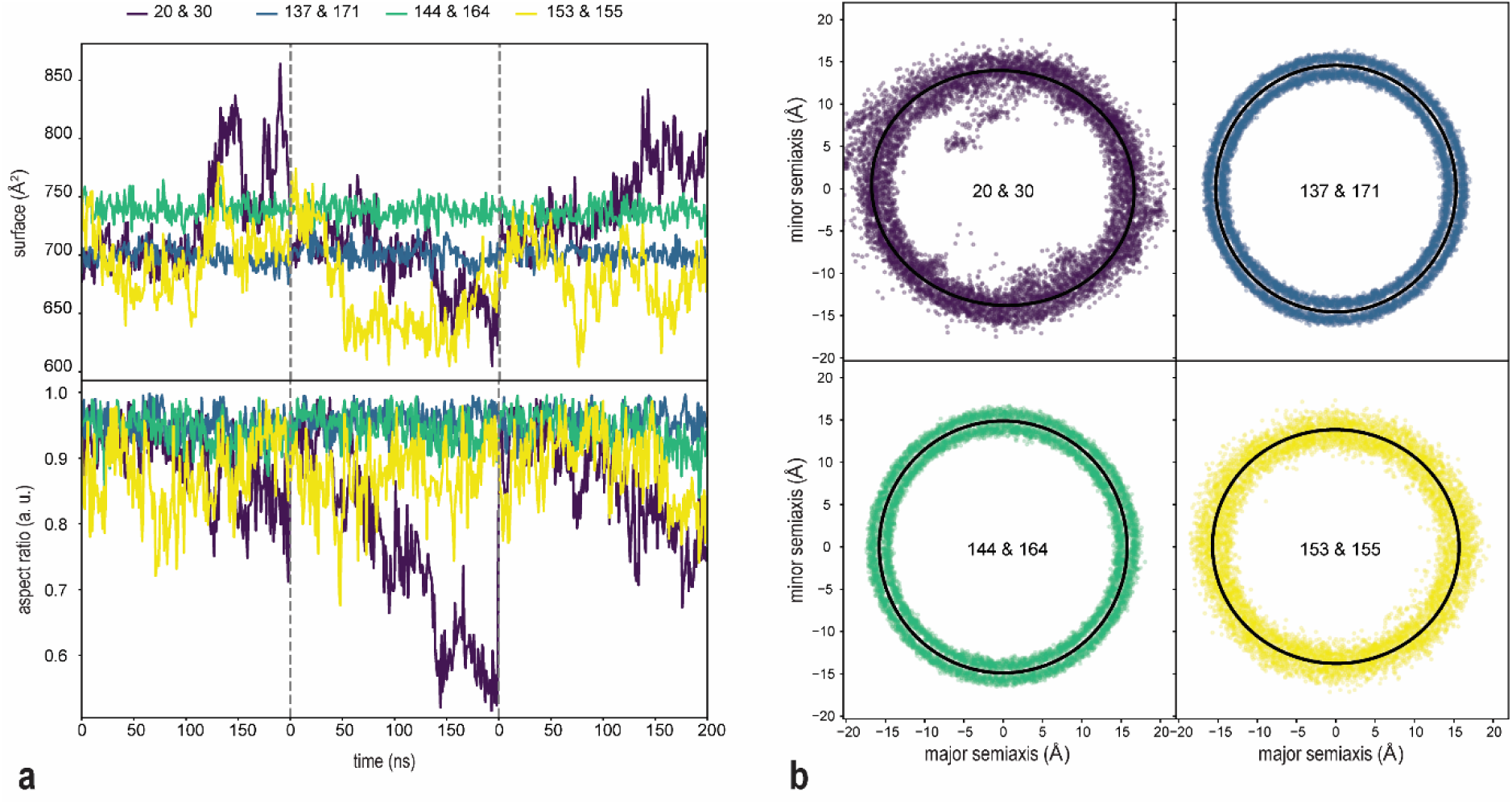
For each conformation in our *C. Perfringens* β-toxin (CPB) simulation, we extracted the coordinates of Cα at each constriction point and fitted them with an ellipse. The constriction point at the intracellular mouth of the toxin (residues 153 and 155) is the most dynamic, featuring the largest fluctuations in the fitted ellipse. (a) time evolution of constriction point surface area and aspect ratio of three consecutive 200ns simulations. (b) position of all extracted Cα coordinates of each constriction point, aligned so that the each fitted ellipse is centered at the origin and oriented so that its major semiaxis is parallel to the x-axis. Black ellipses fitted to these points represent the average constriction points shape. Only the constriction point at the intracellular mouth is noticeably elliptical, with Cα atoms featuring larger deviations from the fitted ellipse.

**Supplemental Figure 8.**
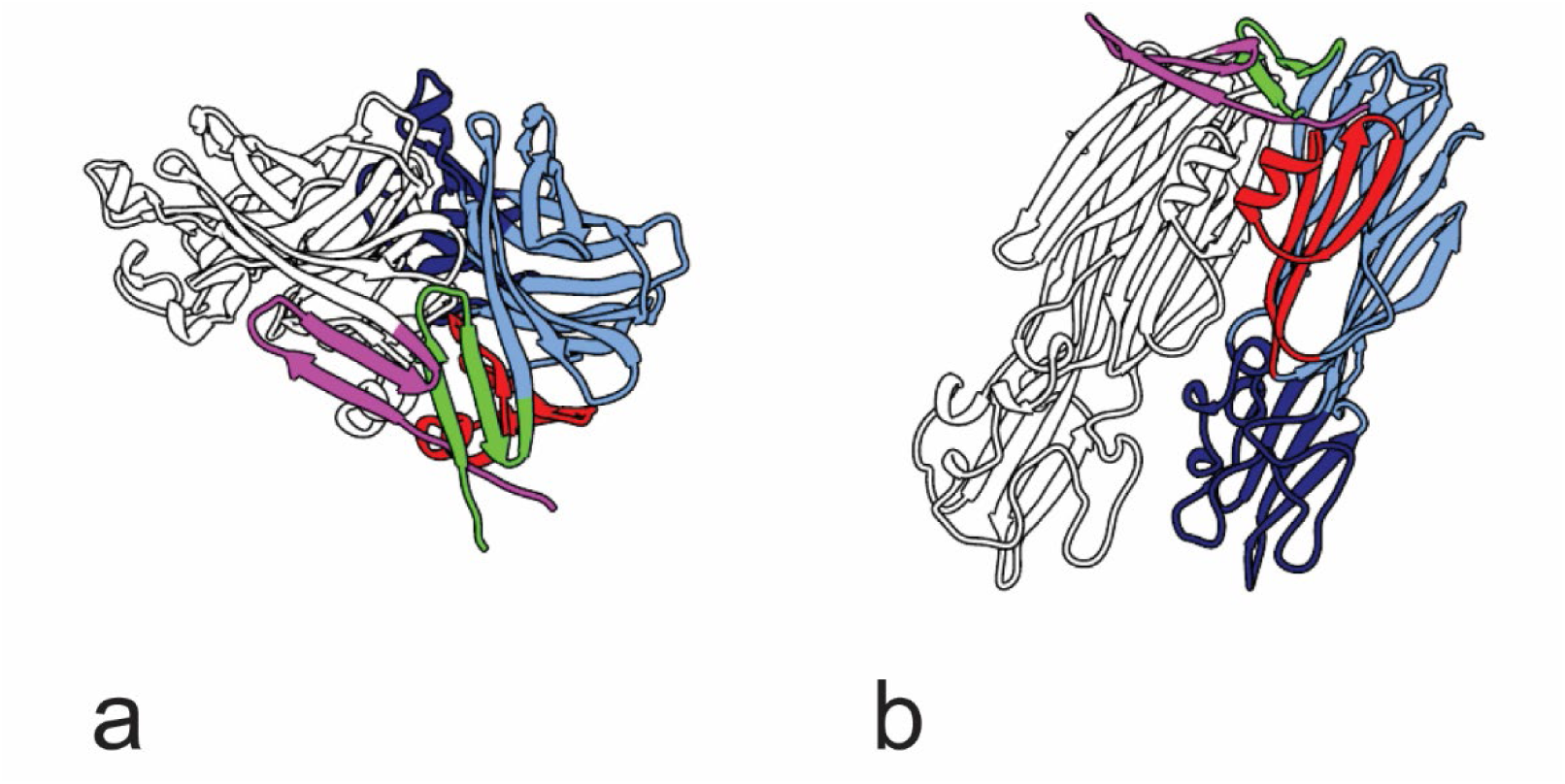
Top (a) and side (b) view of two AlphaFold predicted soluble CPB monomers positioned as consecutive monomers in the CPB oligomer. One monomer is colored as in Fig. 1 while the second is shown in white with the N-terminus in purple highlighting the clash and overlap at the N-terminal region without extracting the NBP from its position during oligomerization.

